# Tomato fruit ripening factor NOR controls leaf senescence

**DOI:** 10.1101/436899

**Authors:** Xuemin Ma, Salma Balazadeh, Bernd Mueller-Roeber

## Abstract

NAC transcription factors (TFs) are important regulators of expressional reprogramming during plant development, stress responses and leaf senescence. NAC TFs also play important roles in fruit ripening. In tomato (*Solanum lycopersicum*), one of the best characterized NAC involved in fruit ripening is NON-RIPENING (NOR) and the *non-ripening* (*nor*) mutation has been widely used to extend fruit shelf life in elite varieties. Here, we show that NOR additionally controls leaf senescence. Expression of *NOR* increases with leaf age, and developmental as well as dark-induced senescence are delayed in the *nor* mutant, while overexpression of *NOR* promotes leaf senescence. Genes associated with chlorophyll degradation as well as senescence-associated genes (SAGs) show reduced and elevated expression, respectively, in *nor* mutants and *NOR* overexpressors. Overexpression of *NOR* also stimulates leaf senescence in *Arabidopsis thaliana*. In tomato, NOR supports senescence by directly and positively regulating the expression of several senescence-associated genes including, besides others, *SlSAG15* and *SlSAG113*, *SlSGR1* and *SlYLS4*. Finally, we find that another senescence control NAC TF, namely SlNAP2, acts upstream of *NOR* to regulate its expression. Our data support a model whereby NAC TFs have often been recruited by higher plants for both, the control of leaf senescence and fruit ripening.

## Introduction

Transcription factors (TFs) of the NAC (for NAM, ATAF1/2 and CUC2) family play important roles for development and the response of plants to abiotic and biotic stresses (Puranik *et al.*, 2012; Shao *et al.*, 2015). A prominent process controlled by NAC TFs is leaf senescence, which is a complex physiological process of nutrient recovery to support the development and growth of newly forming organs, including new leaves, flowers and seeds (Hendelman *et al.*, 2013; Zhong *et al.*, 2016). NAC TFs in diverse dicot and monocot plant species have been shown to control the onset and execution of senescence, e.g. in *Arabidopsis thaliana* (Guo and Gan, 2006; Kim *et al.*, 2009; Balazadeh *et al.*, 2010; Wu *et al.*, 2012; Balazadeh *et al.*, 2014; Garapati *et al.*, 2015; Kamranfar *et al.*, 2018), rice (*Oryza sativum*; Zhou *et al.*, 2013; Mao *et al.*, 2017), wheat (*Triticum aestivum*; Uauy *et al.*, 2006; Zhao *et al.*, 2015), cotton (*Gossypium hirsutum*; Fan *et al.*, 2015), and tomato (*Solanum lycopersicum*; Lira *et al.*, 2017; Ma *et al.*, 2018).

A master positive regulator of leaf senescence in Arabidopsis is ORE1 (ORESARA1; ANAC092; Kim *et al.*, 2009; Balazadeh *et al.*, 2010). Expression of *ORE1* increases with leaf age, a process regulated at the transcriptional level by the *ORE1* promoter, and post-transcriptionally by microRNA *miR164* (Kim *et al.*, 2009). ORE1 controls the expression of a number of senescence-associated genes (SAGs) by directly binding to their promoters (Balazadeh *et al.*, 2010), and accordingly, overexpression or knocking out *ORE1* promotes or inhibits senescence, respectively (Kim *et al.*, 2009; Balazadeh *et al.*, 2010). Recently, the closest putative orthologs of *ORE1* in tomato (i.e. *SlORE1S02*, *SlORE1S03*, and *SlORE1S06*) were also shown to positively control leaf senescence (Lira *et al.*, 2017). In addition, inhibiting *SlORE1S02* by RNA interference (RNAi) not only delayed leaf senescence but also triggered an altered source-sink sugar partitioning resulting in an increased number of fruits per plant with elevated sugar levels (Lira *et al.*, 2017). Similarly, we recently showed that inhibiting expression of the *SlNAP2* transcription factor in transgenic tomato plants delays leaf senescence, which was accompanied by an increased yield of fruits (with elevated sugar content) likely due to extended photosynthesis in aging plants (Ma *et al.*, 2018). *SlNAP2* belongs to the NAP clade of NAC transcription factors of which *AtNAP* from Arabidopsis was first studied with respect to leaf senescence (Guo and Gan, 2006) and was later shown to also control silique senescence (Kou *et al.*, 2012). In rice, inhibiting *OsNAP1* delayed leaf senescence but increased seed yield (Liang *et al.*, 2014).

In addition, NAC TFs have been reported, or suggested, to be involved in ripening of fleshy fruits in several species, with a particular emphasis on tomato, an important fleshy fruit-bearing crop that is extensively used as a model vegetable for studies on fruit physiology and development; its nuclear genome has been sequenced (Tomato Genome Consortium, 2012). One of the best characterized examples in tomato is NON-RIPENING (NOR), which also affects fruit shelf life, an important economic trait. Mutations in the *NOR* gene (locus *Solyc10g006880*) lead to the formation of a truncated TF protein (*nor* mutant) or a NAC TF with a single amino acid substitution (*alcobaca* mutant, *alc*) (Giovannoni *et al.*, 2004; Casals *et al.*, 2012). Recently, a further mutation of the *NOR* gene, leading to an early stop codon, was identified in the tomato variety Penjar-1 grown in the Mediterranean area (Kumar *et al.*, 2018). NOR acts upstream of ethylene synthesis and thereby controls fruit ripening (Barry and Giovannoni, 2007). ChIP assays demonstrated that *NOR* is a direct downstream target of RIN (Ripening Inhibitor), a MADS-box TF controlling fruit ripening (Martel *et al.*, 2011; Fujisawa *et al.*, 2013). Similarly, in melon (*Cucumis melo*), a NOR transcription factor (CmNAC-NOR) was found to be involved in fruit ripening (Ríos *et al.*, 2017). In addition, *NOR* homologs control senescence in non-flesh fruits like the siliques of Arabidopsis where *NARS1*/*NAC2* and *NARS2*/*NAM* redundantly and positively regulate silique senescence while leaf senescence is unaltered compared to wild type, indicating organ-specific functions of the two NAC TFs (Kunieda *et al.*, 2008).

Besides NOR, other TFs of the NAC family in tomato have been reported to control fruit ripening, including SlNAC4 which positively regulates ripening, possibly through physical interaction with NOR and RIN (shown by yeast two-hybrid studies); furthermore, SlNAC4 was suggested to act as an upstream regulator or *RIN* (Zhu *et al.*, 2014). Evidence for a positive role in regulating fruit ripening was also obtained for SlNAC48 and SlNAC19 (which is identical to SlNAP2) using a virus-induced gene silencing (VIGS) approach. The data suggest that both TFs SLNAC47 and SlNAC48 act by affecting ethylene biosynthesis and signaling (Kou et al., 2016). *SlNAC3* shows high expression in fruits and is involved in seed development (Han *et al.*, 2012; Han *et al.*, 2014).

Evidence for an involvement of NAC TFs in fleshy fruit ripening was also obtained from studies performed on developing and ripening fruits of different other species, including the octoploid strawberry cultivar *Fragaria x ananassa* (Moyano *et al.*, 2018), the Chilean endemic strawberry *Fragaria chiloensis* (Carrasco-Orellana *et al.*, 2018), and bilberry (*Vaccinium myrtillus*; Nguyen *et al.*, 2018).

Taken together, many NAC TFs have been reported to control leaf senescence in different plant species, and some NACs have been firmly proven - or suggested - to control the ripening of fleshy or dry fruits. Considering this, we were interested to investigate whether the so-far best studied fruit ripening control NAC TF in tomato, namely NOR, additionally controls leaf senescence in this plant. Our data show that NOR acts as a positive transcriptional regulator of leaf senescence by directly and positively controlling the expression of several chlorophyll degradation-(CDGs) and senescence-associated genes (SAGs) in this species. The data suggest an evolutionary recruitment of NAC TFs from regulating leaf senescence towards the control of physiology during fruit ripening.

## Materials and methods

### General

Tomato orthologs of Arabidopsis genes were identified using the PLAZA 3.0 database (http://bioinformatics.psb.ugent.be/plaza/; Proost *et al.*, 2015). Genes were annotated using the PLAZA 3.0 and Sol Genomics (https://solgenomics.net/) databases, and using information extracted from the literature. Oligonucleotide sequences are given in **Table S1**. qRT-PCR primers were designed using QuantPrime (www.quantprime.de; Arvidsson *et al.*, 2008).

### Plant material and growth conditions

Tomato (*Solanum lycopersicum* L., cultivar Moneymaker) was used as the wild type (WT). The *nor* mutant is in the Rutgers genetic background (Tomato Genetics Research Center, accession LA3013). Seeds were germinated on full-strength Murashige-Skoog (MS) medium containing 2% (w/v) sucrose and 3-week-old seedlings were transferred to soil containing a mixture of potting soil and quartz sand (2:1, v/v). Plants were grown in a growth chamber at 500 μmol photons m^−2^ s^−1^ and 25°C under a 14/10-h light/dark regime in individual pots (18 cm diameter). For experiments with *Arabidopsis thaliana* (L.) Heynh., accession Col-0 was used as the control. Seeds were germinated in soil (Einheitserde GS90; Gebrüder Patzer, Sinntal-Altengronau, Germany) in a climate-controlled chamber with a 16-h day length provided by fluorescent light at approximately 100 μmol m^−2^ s^−1^, day/night temperature of 20°C/16°C, and relative humidity of 60%/75%. After 2 weeks, seedlings were transferred to a growth chamber with a 16-h day (80 or 120 μmol m^−2^ s^−1^), day/night temperature of 22°C/16°C, and 60%/75% relative humidity.

### DNA constructs

Primer sequences are listed in **Table S1**. Amplified fragments generated by PCR were sequenced by Eurofins MWG Operon (Ebersberg, Germany). For *35S:NOR-GFP*, the full-length *NOR* open reading frame was amplified without its stop codon. The PCR product was cloned into the pENTR/D-TOPO vector using the pENTR Directional TOPO Cloning kit (Invitrogen). The sequence-verified entry clone was then transferred to the pK7FWG2 vector (Karimi *et al.*, 2002) by LR recombination (Invitrogen). For *NOR-IOE*, the *NOR* coding sequence was cloned into the pER10 vector (Zuo *et al.*, 2002) made GATEWAY-compatible. Constructs were transformed into tomato cv. Moneymaker using *Agrobacterium tumefaciens* GV2260, or into Arabidopsis using *A. tumefaciens* GV3101 (pMP90).

To construct *NOR-CELD*, the DBP-CELD fusion vector pTacLCELD6XHis was used (Xue, 2005). The NOR coding sequence (without stop codon) was amplified by PCR with a sense primer (including an *Nhe*I restriction site) and an antisense primer (including a *Bam*HI restriction site) (**Table S1**). The amplified DNA fragment was first inserted into pCR2.1 (Thermo Fisher Scientific) and then inserted N-terminal of CELD using the *Nhe*I and *Bam*HI cloning sites of pTacLCELD6XHis to create an in-frame fusion.

### Treatments

For estradiol induction, 3-week-old *NOR-IOE* seedlings were incubated in sterile water containing 15 μM estradiol (control treatment: 0.15% [v/v] ethanol). The seedlings were kept on a rotary shaker for 6 hours and then immediately frozen in liquid nitrogen. For dark-induced leaf senescence experiments, detached young leaves from 10-week-old WT and *NOR* transgenic plants were placed on moisturized filter papers in Petri dishes with the adaxial side facing upwards. The plates were kept in darkness at 22°C for two weeks. Filter papers were changes every five days. Gene expression levels were determined by qRT-PCR.

### Gene expression analysis

Total RNA was extracted using Trizol reagent (Life Technologies). Synthesis of complementary DNA and qRT-PCR using SYBR Green were performed as described (Balazadeh *et al.*, 2008). PCR was performed using an ABI PRISM 7900HT sequence detection system (Applied Biosystems). GAPDH (*Solyc04g009030*) served as reference gene for data analysis. Statistical significance was determined using Student’s *t* test.

### DNA-binding site selection

*In vitro* binding site selection was performed using the CELD-fusion method with the pTacNOR-LCELD6xHis construct, employing biotin-labeled double-stranded oligonucleotides (Xue, 2005). The DNA binding activity of NOR-CELD was measured using methylumbelliferyl β-D-cellobioside as substrate (Xue, 2002). DNA binding assays with a biotin-labeled single-stranded oligonucleotide or a biotin-labeled double-stranded oligonucleotide without a target binding site were used as controls.

### Chromatin immunoprecipitation (ChIP)

ChIP-qPCR was performed from leaves of mature *35S:NOR-GFP* plants, and wild type (WT) served as control. ChIP was performed as described (Kaufmann *et al.*, 2010) using anti-GFP antibody to immuneprecipitate protein-DNA complexes. qPCR primers were designed to flank the NOR binding sites within the promoter regions of potential target genes. Primers annealing to a promoter region of *Solyc04g009030* lacking a NOR binding site were used as a negative control. Primers used for qPCR are listed in **Table S1**.

### Chlorophyll measurements

Chlorophyll content was determined using a SPAD analyser (N-tester; Hydro Agri). Alternatively (Figure 4D), frozen leaf powder was suspended in 5 mL 80% (v/v) acetone in water and homogenized for 1 min. Chlorophyll content was determined with a spectrophotometer at 663 and 646 nm as described by Arnon (1949).

### Ion leakage measurement

Membrane damage during senescence was estimated by measuring ion leakage in control and dark-treated leaves of WT and *NOR*-transgenic plants as reported in Thirumalaikumar *et al.* (2018).

### Accession numbers

Sequence data from this article can be found in the GenBank/EMBL data libraries under the following accession numbers: *NOR* (NM_001247723.2); *SlNAC3* (NM_001279348.2); *SlNAP2* (XM_004236996.2); *SlSAG15* (XM_010320381.2); *SlSAG113* (XP_004239911.1); *SlSGR1* (NP_001234723.1); *SlPPH* (XM_004229633.3); *SlPAO* (NP_001234535.2); *SlYLS4* (XM_004245218); *SlERT1B* (NM_001361347); *SlKFB20* (XM_010320257); *SlABCG40* (XP_004247842.1).

## Results

### *NOR* is upregulated during leaf senescence

*NOR* encodes a tomato NAC transcription factor that harbors a conserved, DNA-binding NAM at its N-terminus (Figure 1A). At the protein level, NOR is closely related to SlNAC3 from tomato, and to NARS1 and NARS2 from Arabidopsis (Figure 1B). To test the subcellular localization of NOR we expressed it as a fusion to green fluorescence protein (GFP) in transgenic tomato plants, under the control of the cauliflower mosaic virus (CaMV) *35S* promoter. As shown in Figure 1C, NOR-GFP fusion protein accumulated in nuclei, as expected for a transcription factor. *NOR* is hardly expressed in young leaves, but its expression increased during developmental and dark-induced senescence (Figure 1D, E), indicating a possible function of the tomato transcription factor for regulating leaf senescence.

**Figure 1.**
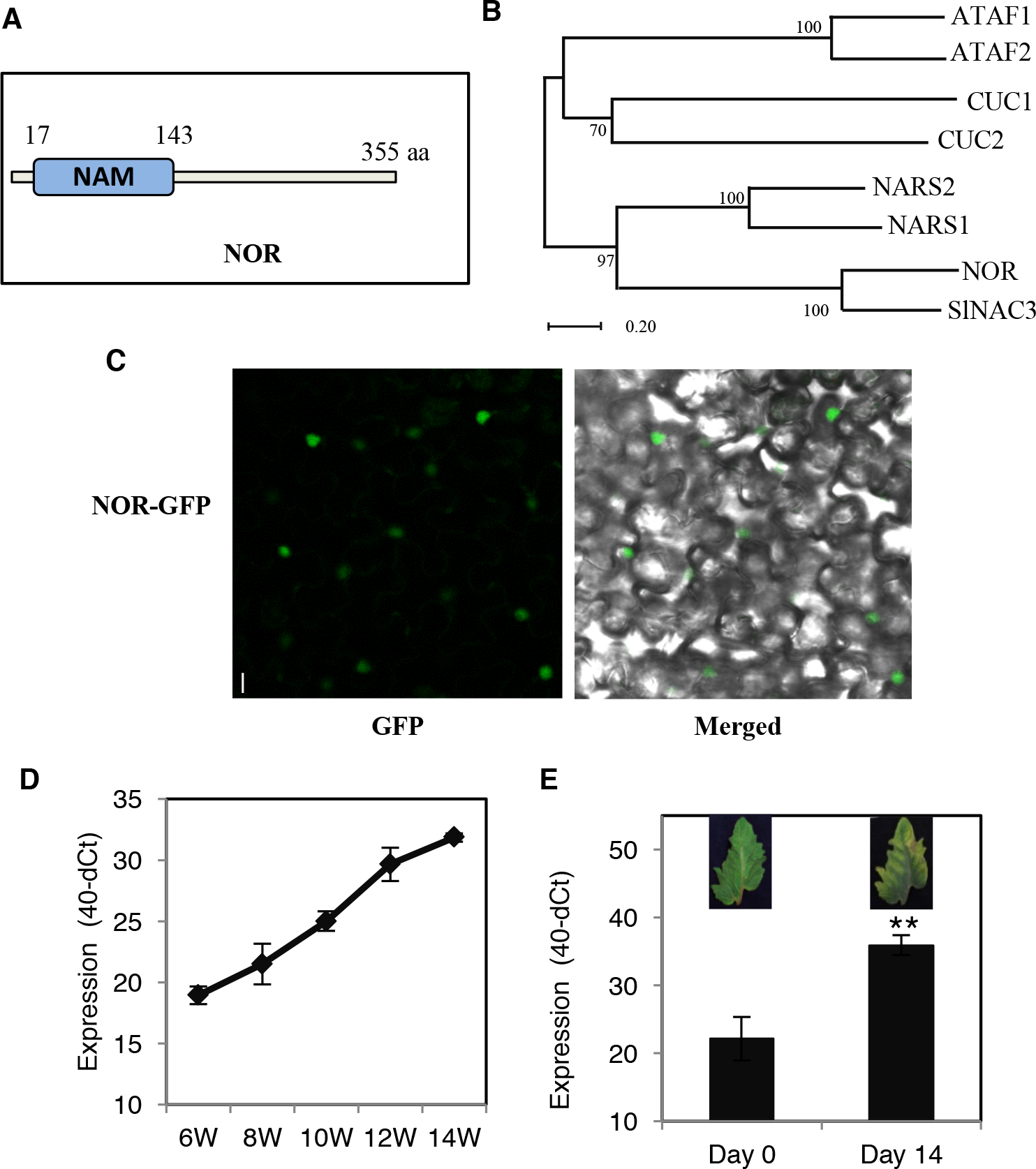
Subcellular localization of NOR and *NOR* expression during senescence. **(A**)Schematic presentation of the NAM domain of NOR. Numbers indicate amino acid positions. **(B**)Phylogenetic analysis of selected NAC proteins. The phylogenetic tree was constructed by MEGA 5.05 software using the neighbor-joining method with the following parameters: bootstrap analysis of 1,000 replicates, Poisson model, and pairwise deletion. NOR and SlNAC3 are two tomato TFs and the others are from Arabidopsis. Gene codes of the Arabidopsis TFs are: *ATAF1*, *At1g01720*; *ATAF2*, *At5g08790*; *NARS1*, *At3g15510*; *NARS2*, *At1g52880*; *CUC1*, *At3g15170*; *CUC2*, *At5g53950.* **(C**)Subcellular localization of NOR protein. NOR fused to GFP was visualized in epidermal cells of transgenic tomato plants by confocal laser scan microscopy. Scale bar, 10 μm. **(D**)*NOR* transcript level in the 3^rd^ true leaf at different developmental stages of tomato wild-type cv. Moneymaker plants. The age of the plants was 6 - 14 weeks (6W - 14W). The y-axis indicates expression level (40- dCt). Data are means ± SD of three biological replicates. **(E**)Expression of *NOR* in young detached leaves of 8-week-old WT plants before (day 0) and after 14 days of dark treatment. Leaves were excised from the top part of the stem. Data are means ± SD (n = 3). Asterisks denote significant difference from Day 0 (Student’s *t*-test, **: *P* ≤ 0.01).

### *NOR* promotes leaf senescence

To test whether NOR indeed regulates leaf senescence, we first generated transgenic tomato (*Solanum lycopersicum* cv. “Moneymaker”) plants constitutively expressing *NOR* under the control of the CaMV *35S* promoter. We selected two lines (hereafter, *OX-L5* and *OX-L19*; **Figure S1A**) for further analysis. Notably, *NOR* overexpression lines showed early leaf senescence, while their stems were also typically shorter than those of wild-type (WT) plants (Figure 2A). The ratio of yellow leaves (defined as leaves with more than 50% yellowing) to all leaves of 12-week-old *OX* plants was significantly higher in *OX-L5* and *OX-L19* plants than the WT (Figure 2B). Furthermore, the chlorophyll content of leaves from the same position (leaf no. 3) dropped faster during development in *OX* than WT (Figure 2C). We also observed a generally reduced shoot height of *NOR* overexpressors compared to the WT, while the *nor* mutant appeared slightly taller under our growth conditions (Figure 2A).

**Figure 2.**
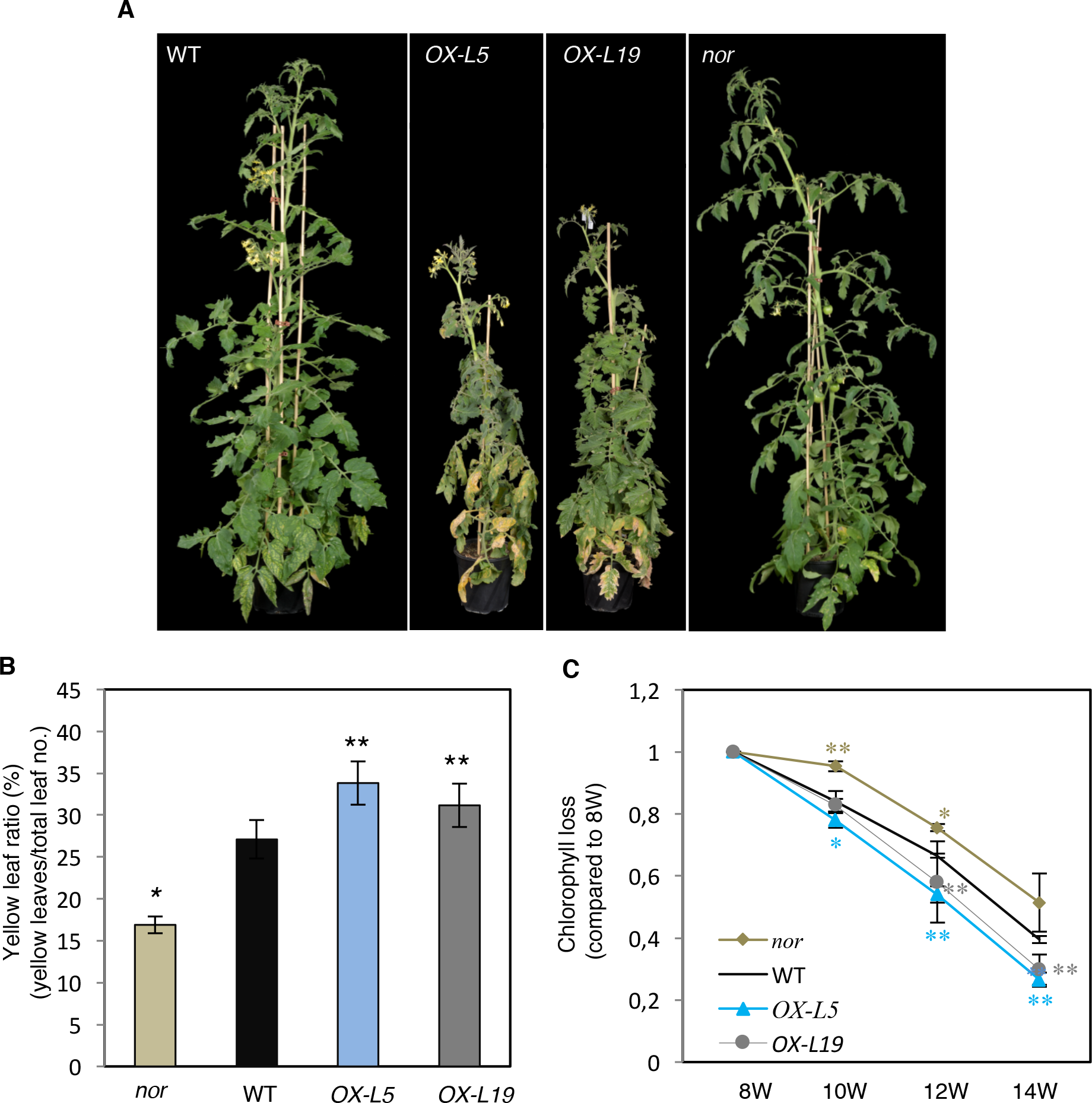
NOR promotes leaf senescence in tomato. **(A**)Phenotype of 12-week-old WT, *OX-L5*, *OX-L19*, and *nor* plants. Note the early leaf senescence in *NOR* overexpressors. **(B**)Yellow leaf ratio of 12-week-old WT, *OX-L5*, *OX-L19*, and *nor* plants. Yellow leaves showing more than 50% yellowing were counted and divided by the total number of leaves. Data are means ± SD (n = 5). **(C**)Chlorophyll loss of the 3^rd^ true leaf (counted from the bottom of the stem) of 8-week- (8W), 10-week- (10W), 12-week- (12W) and 14-week-old (14W) WT, *OX-L5*, *OX-L19*, and *nor* plants. Chlorophyll content was measured by a SPAD meter and at each time point compared to 8W for each genotype (set to 1). Data are means ± SD of three biological replicates. Asterisks in (B) and (C) indicate significant differences from WT (Student’s *t*-test, *: *P* ≤ 0.05; **: *P* ≤ 0.01).

### The tomato *nor* mutants exhibits retarded leaf senescence

Dark treatment is an efficient way to induce senescence in plants, as shown in many reports (Biswal and Mohanty, 1976; Chen and Kao 1991; Weaver *et al.*, 2001). We therefore examined the phenotypes of tomato *nor*, WT, and *OX-L19* plants after 14 days of dark treatment. Detached leaves from the overexpression line showed earlier de-greening in extended darkness than the WT. In contrast, leaves of the *nor* mutant remained longer green in darkness and their chlorophyll content remained high after treatment compared to WT and *OX-L19* (Figure 3A, B). Moreover, ion leakage, an indicator of membrane damage, was significantly elevated in *OX-L19* compared to WT, while it was reduced in *nor* (Figure 3C). In accordance with this, expression of various senescence-associated genes (SAGs) and chlorophyll degradation genes (CDGs) was upregulated in *OX-L19* plants compared to wild type, but downregulated in *nor* (Figure 3D; **Table S2**).

**Figure 3.**
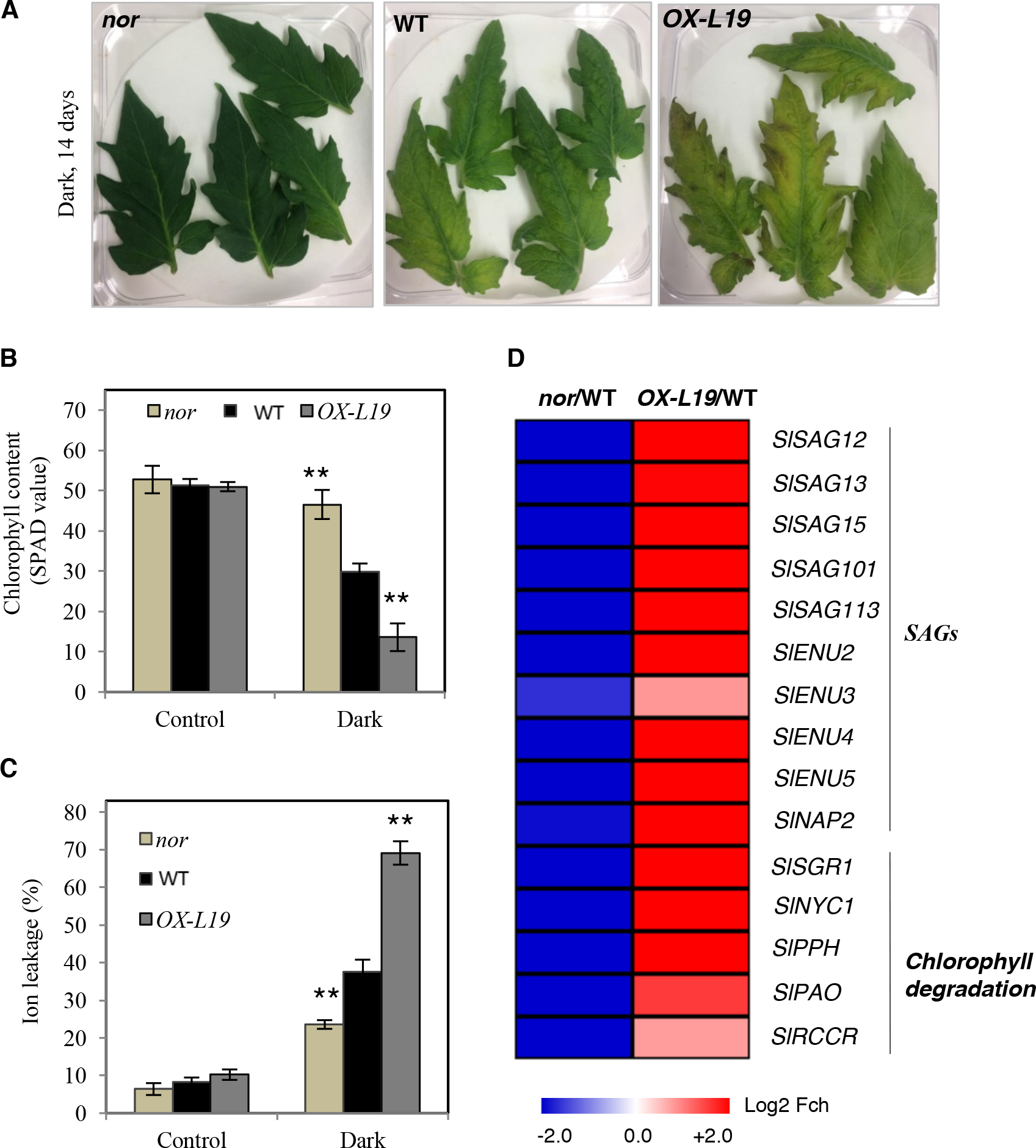
Dark-induced leaf senescence in *NOR*-modified plants. **(A)** Detached leaves of 8-week-old *nor*, WT, and *OX-L19* plants after dark treatment. Young leaves from the top of the stem were detached and subjected to darkness for 14 days (Dark). (B) Chlorophyll content of leaves before darkness (control) and of dark-treated leaves. Chlorophyll content was measured using a SPAD meter. **(C**) Ion leakage of leaves before (control) and after dark treatment. **(D**) Heat map showing the fold change (log_2_) of the expression of *SAG*s and chlorophyll degradation genes in detached leaves of 8-week-old plants *nor* and *OX-L19*, after dark treatment, compared to WT. The full data are given in **Table S2**. In ((B) and (C), asterisks indicate significant differences from the WT (Student’s *t*-test; **: *P* ≤ 0.01).

To further examine the function of NOR in regulating senescence, we generated *NOR* knock-down lines by artificial microRNA (*ami-NOR*), in tomato cultivar Moneymaker. The *ami-NOR* construct targets 21 nucleotides (TGTACCATAGTTTGAAGGCTG) around 200 bp close to 3’ end of the *NOR* coding sequence. This region encodes the transactivation domain of the TF. We selected two lines (*ami-L2* and *ami-L35*) with a reduced *NOR* transcript abundance as determined by end-point PCR (**Figure S2A**). The *ami-NOR* lines exhibited delayed senescence during dark treatment, similar to the *nor* mutant (**Figure S2B and S2C**).

### NOR promotes leaf senescence in Arabidopsis

To test whether *NOR* also induced early leaf senescence in a heterologous species, we overexpressed it in transgenic *Arabidopsis thaliana* plants. We selected two Arabidopsis lines expressing *NOR* for further analysis (hereafter, *OX-L6* and *OX-L8*; Figure 4A). As in tomato, overexpression of *NOR* promoted early leaf senescence in Arabidopsis (Figure 4A), indicating functional conservation across species. *OX* plants had a higher ratio of yellow to all leaves than the WT at the same age (5 weeks) (Figure 4B).

**Figure 4.**
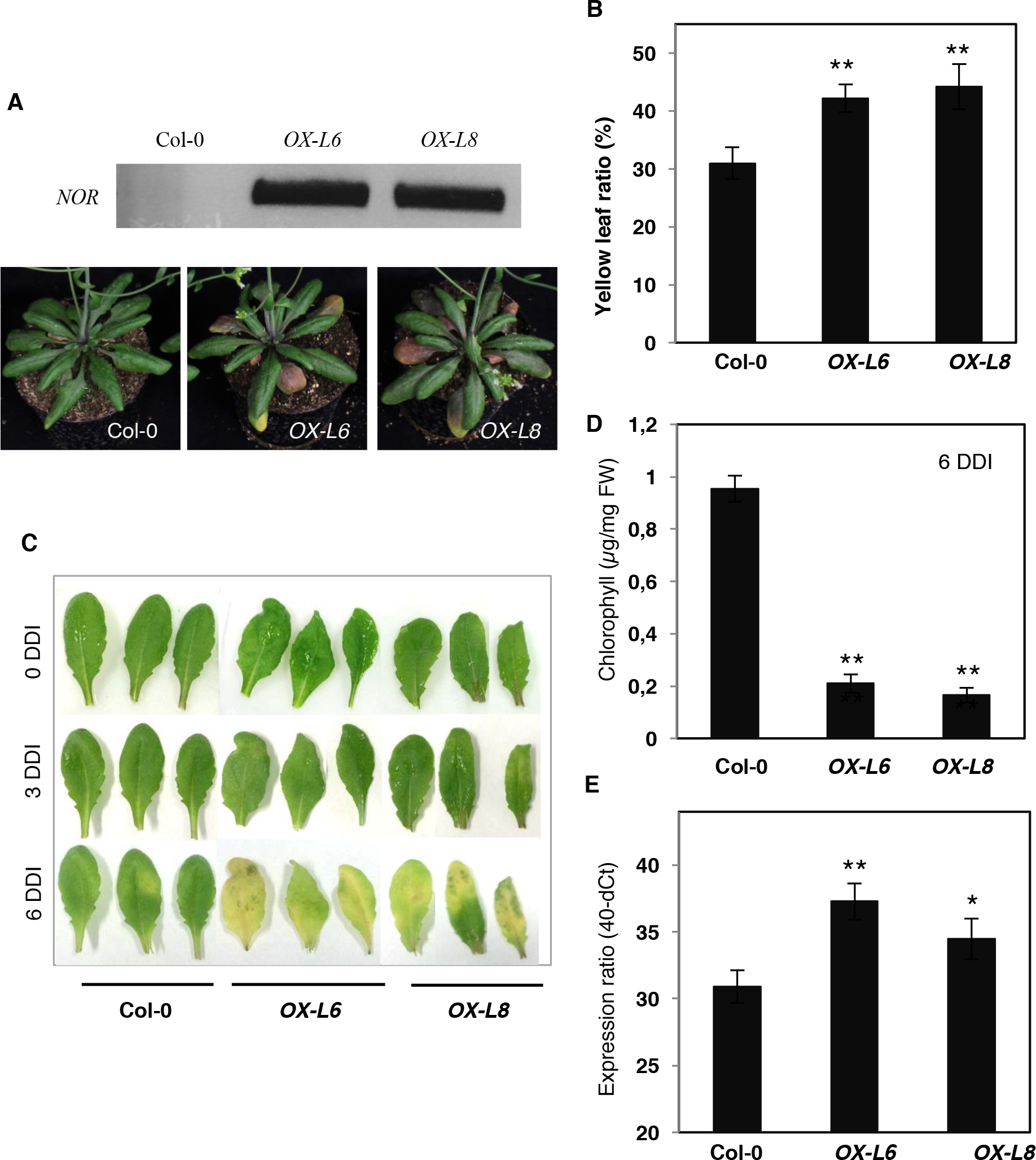
Overexpressing *NOR* in Arabidopsis promotes leaf senescence. **(A**) Phenotype of Arabidopsis Col-0 wild-type and *NOR* overexpression plants. The upper panel shows *NOR* transcript abundance in *OX-L6* and *OX-L8* plants, determined by end-point PCR; as expected, no *NOR* transcript is observed in the Arabidopsis WT. The lower panel shows the phenotype of 5-week-old plants (Col-0 and *NOR* overexpressors). **(B**) Yellow leaf ratio of 5-week-old Col-0, *OX-L6*, and *OX-L8* plants. Yellow leaves showing more than 50% yellowing were counted and compared to the total leaf number. Data are means ± SD (n = 5). **(C**) Dark-induced senescence. DDI, days after dark incubation. Note the more pronounced senescence in the two *NOR* overexpressors compared to Col-0 at 6 DDI. Leaves no. 5 - 7 were detached from the various plants were used in the experiment. **(D**) Chlorophyll content of (C), at 6 DDI of Col-0, *OX-L6*, and *OX-L8* plants (n = 5). **(E**) Expression of *AtSAG12* in detached leaves no. 5 - 7 of Col-0, *OX-L6*, and *OX-L8* plants at 6 DDI. The y-axis indicates expression level (40-dCt). Data are means ± SD of three biological replicates. Asterisks in (B), (D) and (E) indicate significant difference from the Col-0 wild type (Student’s *t*-test; *: *P*≤ 0.05; **: *P* ≤ 0.01).

To test whether *NOR* overexpression promotes senescence in darkness, we detached leaves from the Arabidopsis *OX-L6* and *OX-L8* lines and after 6 days of dark incubation observed much stronger senescence than in leaves of the WT control (Figure 4C). Chlorophyll content after dark treatment was more strongly reduced in these lines than the WT (Figure 4D). Expression of the senescence-associated marker gene *AtSAG12* (Noh and Amasino, 1999) was significantly upregulated in these lines in comparison to WT (Figure 4E). From these results, we conclude that NOR positively regulates leaf senescence in both, tomato and Arabidopsis.

### Identification of the consensus DNA binding sequence of NOR

Knowledge about the DNA binding motif(s) of a TF under analysis strongly assists in unraveling the wider gene regulatory network it controls. We therefore performed an *in vitro* binding site selection assay using the earlier reported cellulose D (CELD) fusion method (Xue, 2005) to identify NOR binding sites. We first analyzed the binding activity of NOR toward 16 randomly selected TaNAC69 motifs, S1 - S16, bound by the NAC69 transcription factor from wheat (*Triticum aestivum*) (**Figure S3A**). Previously, it was shown that S1 is a high-affinity binding sequence of TaNAC69 (Xue *et al.*, 2006). In our results, NOR showed strong binding affinity to S1, with affinity decreasing progressively with substitutions. Overall, NOR bound to TaNAC69-selected motifs containing the YACG (or CGTR) core sequence (**Figure S3A**). Further analysis of the specificity of binding through base substitution, insertion, or deletion revealed that the mutation of nucleotides in the core motifs (e.g., S1m3 and S1m9) resulted in a strong reduction of NOR binding activity (**Figure S3B**). Taken together, our data suggest two high-affinity binding sites of NOR, CG(Y/C)(G/C)(5-7n)N(A/G)CGn(A/C/G)(A/C/T) and (C/T)ACGn(A/C)(A/T)(C/G/T)(C/T), as motif I and motif II, respectively.

### Identification of NOR target genes

Although NOR is a transcription factor well known for its function in fruit ripening, no direct target genes have to our knowledge been reported so far. Therefore, based on the results presented in Figure 3D, we selected individual genes for further analysis to test whether they might be direct downstream targets of NOR. To this end, we chose several genes harboring the NOR binding site within their 5´ upstream regulatory regions, including *SlSAG15*, *SlSAG113*, *SlSGR1*, *SlPPH*, and *SlPAO* (Figure 5A). Chromatin-immunoprecipitation/quantitative real-time PCR (ChIP-qPCR) revealed direct binding of the NOR transcription factor to the promoters of all genes except *SlPAO* (Figure 5B).

**Figure 5.**
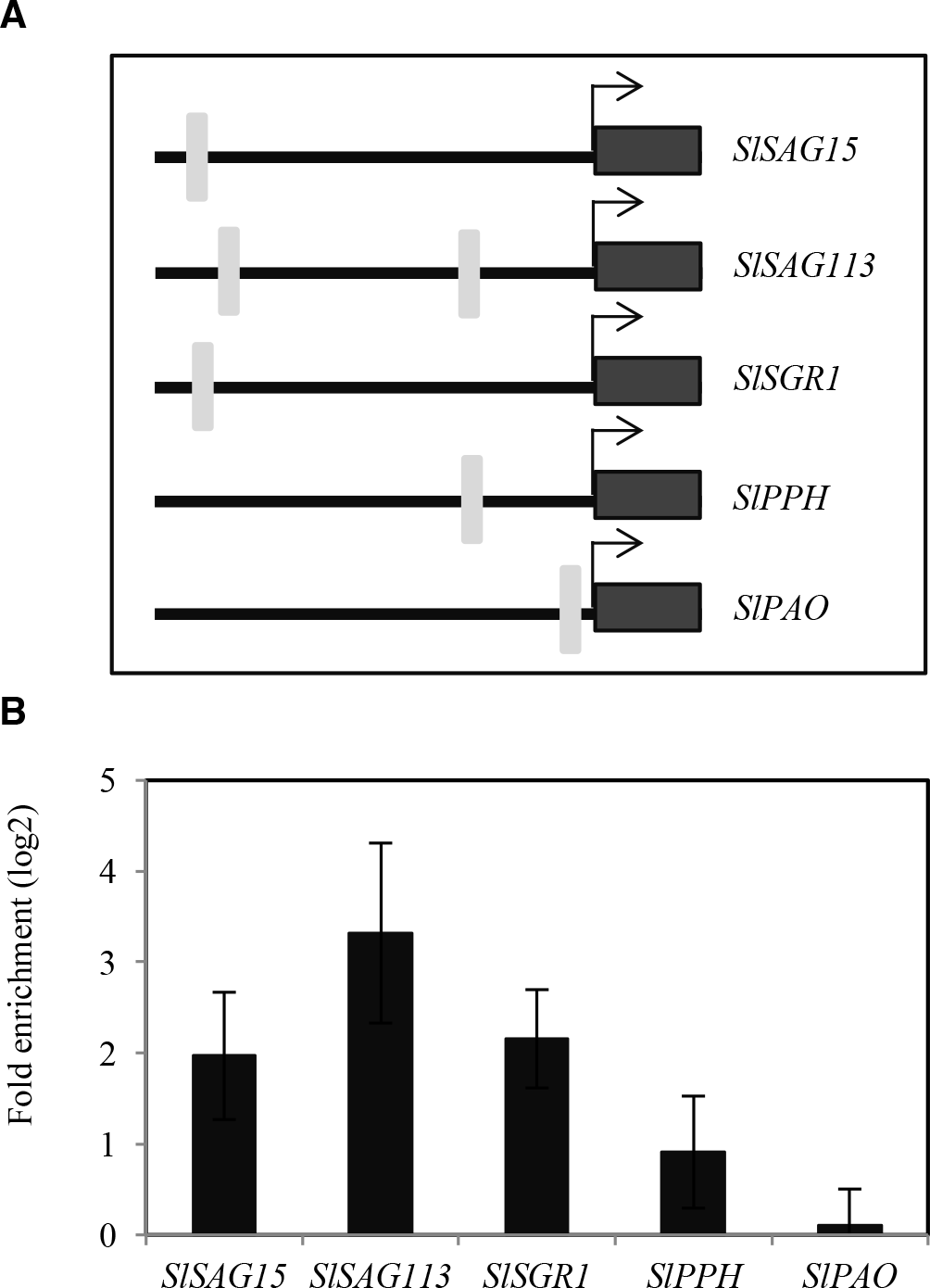
Direct regulation of SAGs by NOR. **(A**)Schematic diagram showing positions of NOR binding sites in 1-kb promoters of selected genes. Arrows indicates the ATG translational start codon. Light-grey boxes indicate the NOR binding sites and black boxes indicate the coding regions of the genes. Sequences of the gene promoters including the NOR binding sites tested in the ChIP experiments are given in **Table S3**. **(B**)ChIP-qPCR shows enrichment of *SlSAG15, SlSAG113*, *SISGR1* and *SlPPH* promoter (1 kb) regions containing the NOR binding site. Eight-week-old *NOR-GFP* plants (mature leaves no. ~3‒5) were harvested for the ChIP experiment. qPCR was performed to quantify the enrichment of the promoter regions. In the case of *SlSAG113*, which has two potential NOR binding sites in its promoter (see panel A), we tested binding of NOR to the sequence proximal to the *ATG* start codon. Values were normalized to the values for *Solyc04G009030* (promoter lacking a NOR binding site). Data are the means ± SD of two independent biological replicates, each determined in three technical replicates.

We next selected additional genes known to be regulated by natural or dark-induced senescence in tomato, or induced by abiotic stresses that trigger senescence (based on literature reports) and checked whether their promoters harbor a NOR binding site. Considering that NOR regulates fruit ripening (Giovannoni *et al.*, 2004; Casals *et al.*, 2012; Kumar *et al.*, 2018) we also included a few genes reported to control this process. We then tested whether expression of these genes is affected in transgenic tomato plants expressing the NOR transcription factor under the control of an estradiol (EST)-inducible promoter (hereafter, *NOR-IOE*). As shown in **Figure S1B**, expression of *NOR* was strongly enhanced in three-week-old *NOR-IOE* seedlings 6 hours after treatment with 15 μM EST, as expected. Similarly, all selected NOR-binding site-containing genes except two showed enhanced expression when *NOR* was induced (Figure 6A; **Table S2**).

**Figure 6.**
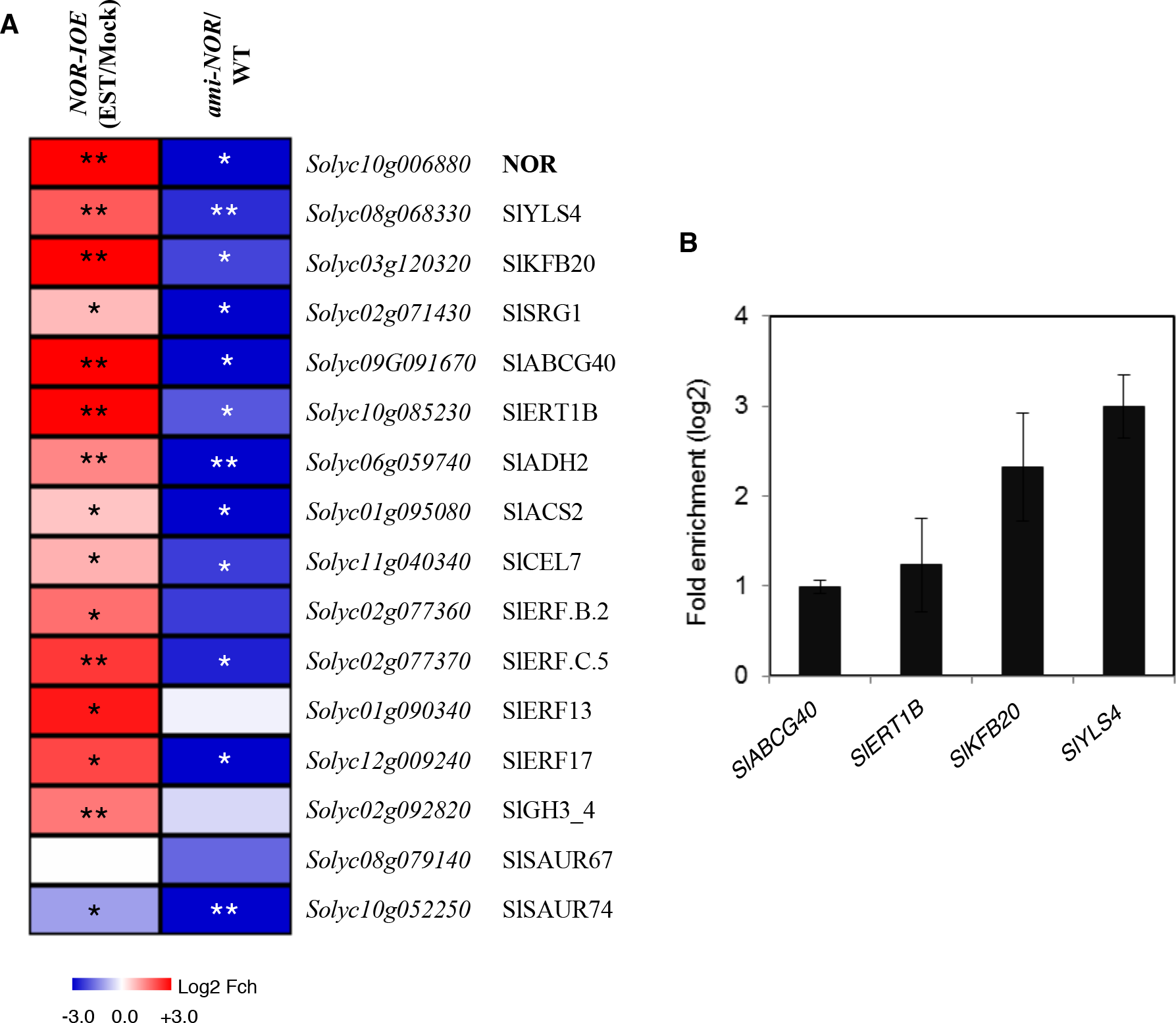
Heat map of differentially expressed genes in *NOR-IOE* and *ami-NOR* plants. **(A**)Gene expression was analyzed by qRT-PCR in *NOR-IOE* seedlings treated with EST (15μM) for 6 h and compared to expression in mock-treated (ethanol, 0.15% [v/v]) seedlings (left column), or in *ami-NOR* seedlings compared to wild-type (WT) seedlings. Seedlings were three weeks old. The color code indicates the log_2_ scale of the fold change; blue, downregulated; red, upregulated. Data represent means of three biological replicates. Data are means ± SD of three biological replicates. Asterisks indicate significant difference from mock-treated samples (for *NOR-IOE* samples) or from WT (for *ami-NOR* samples). Student’s *t*-test; *: *P* ≤ 0.05; **: *P* ≤ 0.01). The full data are given in **Table S2**. **(B**)ChIP‒qPCR shows enrichment of *SlABCG40*, *SlERT1B*, *SlKFB20*, and *SlYLS4* promoter regions containing the NOR binding site within the 1-kb upstream promoter regions of the corresponding genes. Experimental conditions were as described in legend to Figure 5B. Sequences of the gene promoters including the NOR binding sites tested in the ChIP experiments are given in **Table S3**. Data are the means ± SD of two independent biological replicates, each determined in three technical replicates.

Among the genes upregulated by NOR are the senescence-related genes *SlYLS4*, *SlKFB20*, and *SlSRG1*. *SlYSL4* (*Solyc08g068330*), a homolog of Arabidopsis *YLS4* (*YELLOW LEAF SPECIFIC4*), is expressed in a senescence-specific manner; the gene encodes an aspartate aminotransferase possibly involved in remobilizing leaf nitrogen during senescence (Yoshida *et al.*, 2001). *SlKFB20* (*Solyc03g120320*), a gene induced in tomato leaves during senescence, is a homolog of Arabidopsis *AT1G80440*, which encodes a kelch-repeat F-box protein targeting type-B ARR (Arabidopsis Response Regulator) proteins for degradation in the negative regulation of the cytokinin response (Kim *et al.*, 2013a; Kim *et al.*, 2013b). Notably, cytokinins delay senescence (Hwang *et al.*, 2012). *SlSRG1* (*Solyc02g071430*) is closely related to *SENESCENCE-RELATED GENE1* (*SRG1*) from Arabidopsis, which encodes a member of the Fe (II)/ascorbate oxidase gene family and is highly induced at low-nitrogen condition and during sucrose-induced senescence (Pourtau *et al.*, 2006). *SlABCG40* (*Solyc09g091670*), which encodes a protein belonging to the ATP binding cassette (ABC) transporters, is one of the most upregulated genes after EST treatment. It is induced by more than 120-fold after induction of *NOR* with EST in *NOR-IOE* lines. In Arabidopsis, *ABCG40* encodes an ATP binding cassette (ABC) transporter protein involved in the cellular uptake of abscisic acid (ABA; Kang *et al.*, 2010), a phytohormone that triggers stomatal closure upon water shortage and stimulates leaf senescence in various species (Zhang *et al.*, 2012; Zhao *et al.*, 2017).

Three other genes analyzed, namely *SlERT1B*, *SlADH2*, and *SlACS2*, are involved in fruit ripening, and all are upregulated after EST treatment in *NOR-IOE* plants. *SlERT1B* (*Solyc10g085230*), encodes a putative UDP-glycosyltransferase potentially involved in glycoalkaloid biosynthesis in tomato fruits (Itkin *et al.*, 2013; Alseekh *et al.*, 2015). *SlADH2* (*ALCOHOL DEHYDROGENASE2*; *Solyc06g059740*) participates in the biosynthesis of volatiles and, accordingly, its transcript abundance increases during fruit ripening (Speirs *et al.*, 1998); it is a direct target of RIN (Qin *et al.*, 2012). *SlACS2* (*1-AMINOCYCLOPROPANE*- *1*-*CARBOXYLATE SYNTHASE2*; *Solyc01g095080*) encodes an ethylene biosynthesis gene highly expressed during fruit ripening. Downregulation of *SlACS2* lowers ethylene production and delays fruit ripening (Oeller *et al.*, 1991). In addition, expression of *SlACS2* is largely dependent on transcription factor RIN, which is a direct upstream regulator of it (Martel *et al.*, 2011).

We included further genes with likely functions in fruit ripening or leaf senescence in our analysis. One is *SlCEL7* (*Solyc11g040340*), which encodes a putative endo-β-1,4-glucanase of the glycosyl hydrolase 9 (cellulase E) family (www.uniprot.org); *SlCEL7* has been suggested to play a specific role for regulating the loosening of cells walls during fruit growth (Catalá *et al.*, 2000). As seen in Figure 6A(and **Table S2**), expression of *SlCEL7* was significantly elevated in *NOR-IOE* plants after EST induction, suggesting it to be a downstream target of NOR. In addition, the hormone-related genes *ETHYLENE-RESPONSIVE TRANSCRIPTION FACTOR B.2* (*SlERF.B.2*, *Solyc02g077360*), *SlERF.C.5* (*Solyc02g077370*), *SlERF13* (*Solyc01g090340*) and *SlERF17* (*Solyc12g009240*) were also significantly upregulated after EST induction of the NOR transcription factor (Figure 6A), suggesting them to be downstream targets of NOR.

As the phytohormone auxin is involved in controlling leaf senescence and fruit ripening (Kim *et al.*, 2011; Breitel *et al.*, 2016), we also included three auxin-related genes in our analysis, namely *SlGH3_4* (*Solyc02g092820*), which encodes a putative indole-3-acetic acid amido synthetase, an enzyme conjugating auxin to an inactive form thereby reducing cellular free auxin levels, and small auxin up-regulated RNA67 (*SlSAUR67*; *Solyc08g079140*). Expression of various *GH3* genes has previously been shown to increase in leaves during developmental and dark-induced senescence, consistent with the decrease of free auxin levels in senescing leaves (Buchanan-Wollaston *et al.*, 2005; van der Graaff *et al.*, 2006; Kim *et al.*, 2011). While *SlGH3_4* was significantly upregulated upon induction of *NOR*, *SlSAUR67* was not affected. Expression of *SlSAUR74* (*Solyc10g052550*) was significantly reduced after *NOR* induction (Figure 6A).

We next analyzed expression of the selected genes by qRT-PCR in *ami-NOR* lines. Almost all genes that were upregulated in *NOR-IOE* plants after EST induction were downregulated in *ami-NOR* confirming the transcription activation role of NOR toward these genes (Figure 6A).

Finally, we employed ChIP-qPCR to test binding of NOR to the promoters of selected downstream targets *in vivo*, including *SlABCG40*, *SlERT1B*, *SlKFB20* and *SlYLS4.* As shown in Figure 6B, NOR binds to all four promoters.

### SlNAP2 affects *NOR* expression

We previously reported that SlNAP2, a tomato NAC transcription factor, functions as a positive regulator of leaf senescence by directly controlling the expression of various senescence-associated genes as direct targets. In addition, SlNAP2 controls the expression of several ABA-related genes (Ma *et al.*, 2018). SlNAP2 has two related DNA-binding sites, called BS1 and BS2, which are present in the promoters of its direct gene targets (Ma *et al.*, 2018). As previous work on Arabidopsis indicated regulatory connectivity between different NAC TFs to control senescence (e.g. Garapati *et al.*, 2015; Kim *et al.*, 2018), we here thought to investigate the possibility that *NOR* is a downstream affected gene target of SlNAP2. In accordance with this model, sequence analysis of the *NOR* promoter identified an SlNAP2 BS1 binding site 403 bp upstream of the ATG start codon (Figure 7A). Furthermore, expression of *NOR* significantly increased in transgenic tomato plants expressing *SlNAP2* from an EST-inducible promoter (*SlNAP2-IOE*; Ma *et al.*, 2018) 6 h after EST treatment (Figure 7B). Finally, SlNAP2 directly binds to the *NOR* promoter, as shown by ChIP-qPCR (Figure 7C). Collectively, our data thus show that SlNAP2 functions an upstream regulator of *NOR.*

**Figure 7.**
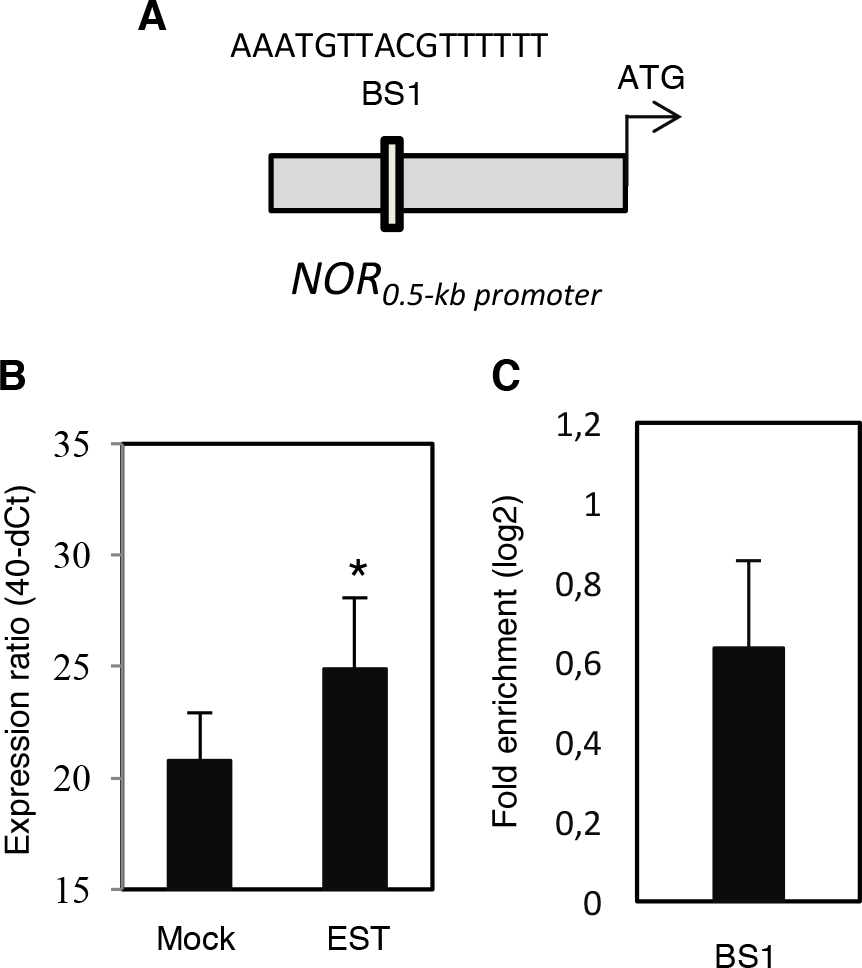
SlNAP2 acts as an upstream regulator of *NOR.* **(A**) Schematic presentation of the SlNAP2 binding site 1 (BS1) within the *NOR* promoter. The sequence of the binding site, which is located in the forward strand of the promoter, is indicted. **(B**) Expression of *NOR* in in 3-week-old *SlNAP2-IOE* seedlings treated with estradiol (EST; 15 μM) for 6 h compared to ethanol (0.15% [v/v])-treated seedlings (Mock). Gene expression was determined by qRT-PCR. Data represent means of three biological replicates. Asterisks indicate significant difference from mock-treated plants (Student’s *t*-test; *: P ≤ 0.05). **(C**) ChIP-qPCR shows enrichment of the *NOR* promoter region containing the SINAP2 binding site 1 (BS1). Mature leaves (no. 3 - 5) harvested from 8-week-old *SlNAP2-GPF* plants were used for the ChIP experiment. Values were normalized to the values for *Solyc04g009030* (promoter lacking a SlNAP2 binding site). Data are means ± SD of two independent biological replicates, each performed with three technical replicates.

## Discussion

NOR-RIPENING (NOR) is a NAC transcription factor well characterized for its role in fruit ripening in tomato (Barry and Giovannoni, 2007; Casals *et al.*, 2012; Kumar *et al.*, 2018). Also in melon (*Cucumis melo*) a NOR homologue has been shown to affect fruit ripening (Ríos *et al.*, 2017). Recently, several other NAC TFs have been reported to control fruit ripening in tomato, including e.g. SlNAC4 (Zhu *et al.*, 2014), SlNAC19 and SlNAC48 (Kou *et al.*, 2016), while SlNAC3 has a function in seed development (Han *et al.*, 2012; Han *et al.*, 2014). With the data available so far, it appears that NAC TFs - in conjunction with other TFs of other families - form interconnected regulatory networks to control fruit aging. For example, RIN, a long-known regulator of tomato fruit ripening of the MADS-box TF family, directly regulates *NOR* by binding to its promoter, as revealed by ChIP assay (Martel *et al.*, 2011; Fujisawa *et al.*, 2013). In addition, expression of *NOR* and *RIN* is reduced in *SlNAC4* RNA interference lines which might indicate that it acts as an upstream regulator of *NOR* and *RIN* (Zhu *et al.*, 2014). Furthermore, yeast two-hybrid assays revealed an interaction of SlNAC4 protein with NOR and RIN, although a functional relevance of this interaction *in planta* was not demonstrated (Zhu *et al.*, 2014). Recently, experimental evidence showed that the basic leucine zipper (bZIP) transcription factor SlAREB1, which at the transcript level is induced by ABA, may function as an upstream regulator of *NOR*, although direct *in planta* binding of SlAREB1 to the *NOR* promoter by e.g. chromatin-immunoprecipitation (ChIP) was not demonstrated (Mou *et al.*, 2018).

Although increasing evidence suggests an involvement of multiple NAC factors in tomato fruit development, a role of NACs in the regulation of leaf senescence in this vegetable crop has rarely been demonstrated despite the fact that NACs play diverse functions in the control of leaf senescence in other species (Podzimska-Sroka *et al.*, 2015; Leng *et al.*, 2017; Ma *et al.*, 2018; Yang and Udvardi, 2018). A particular detailed knowledge about the NAC-controlled senescence networks is available for Arabidopsis where multiple NAC TFs have been shown to positively or negatively regulate leaf senescence by binding to the promoters of diverse target genes to control different physiological processes underlying the complex syndrome of senescence (Guo and Gan, 2006; Kim *et al.*, 2009; Wu *et al.*, 2012; Garapati *et al.*, 2015; Sakuraba *et al.*, 2016; Oda-Yamamizo *et al.*, 2016; Kamranfar *et al.*, 2018; Kim *et al.*, 2018; Li *et al.*, 2018).

The situation is less clear in tomato, although aging in fruits and leaves may at least in part share identical TFs and gene regulatory networks. Recently, Lira *et al.* (2017) found that orthologs of *ORE1*, a central positive regulator of leaf senescence in Arabidopsis (Kim *et al.*, 2009; Balazadeh *et al.*, 2010), control leaf senescence in tomato leading to extended greenness upon downregulation of *SlORE1* gene expression. The increased fruit yield in such plants might be due to an extended photosynthetic lifetime of the leaves, providing carbon for fruit (sink) growth over a longer period than in wild-type plants, although another possibility is that *SlORE1* genes directly control ripening processes in fruits. Similarly, downregulation of NAC transcription factor *SlNAP2* expression delays leaf senescence in tomato followed by an increased fruit yield (Ma *et al.*, 2018). SlNAP2 binds to the promoters of several senescence-related genes, including *SlSAG113* (*Solanum lycopersicum SENESCENCE-ASSOCIATED GENE113*) and the chlorophyll-degradation genes *SlSGR1* (*S. lycopersicum senescence-inducible chloroplast stay-green protein 1*) and *SlPAO* (*S. lycopersicum pheide a oxygenase*). SlNAP2 also directly controls the expression of several abscisic acid (ABA)-related genes including ABA transport, biosynthesis and degradation genes, suggesting that it has an important function in controlling ABA homeostasis in senescing tomato leaves (Ma *et al.*, 2018).

Here, we report that the long-known tomato fruit ripening factor NOR controls leaf senescence, thereby identifying a novel role of NOR for controlling development. Of note, overexpression of *NOR* in both, transgenic tomato and Arabidopsis plants promotes developmental leaf senescence as well as dark-induced senescence. The role of NOR in regulating leaf senescence is related to changes in the expression of senescence-associated genes (SAGs) and chlorophyll degradation genes (CDGs). Expression of various senescence-related genes was enhanced in constitutive or estradiol-inducible *NOR* overexpressors, and we demonstrated binding of the NOR TF to their promoters by chromatin-immunoprecipitation (ChIP). As direct *in vivo* targets of NOR we identified *SlSAG15*, *SlSAG113*, *SlSGR1* and *SlPPH*, as well *SlABCG40*, *SlERT1B*, *SlKFB20* and *SlYLS4.* Of note, *SlSAG113*, *SlSGR1*, and *SlABCG40* were previously also identified as direct targets of SlNAP2 (Ma *et al.*, 2018), strongly suggesting functional overlap of NOR and SlNAP2 in regulating leaf senescence-associated genes in tomato. In accordance with this, both TFs belong to the same clade (the NAP clade) of NAC factors (Kou *et al.*, 2014). This clade also includes the *AtNAP* gene, a well-known regulator of leaf senescence in *A. thaliana* (Guo and Gan, 2006). Interestingly, however, *SlPAO* did not appear to be a direct downstream target gene of NOR (this report; Figure 5B), while we previously found it to be a direct target of SlNAP2 (Ma *et al.*, 2018), indicating partial, but not complete, functional redundancy of both TFs with respect to the control of leaf senescence. Such a redundancy of NAC TFs for the control of senescence was recently highlighted for Arabidopsis by Li *et al.* (2018).

Another important finding of our study is that *SlNAP2* itself is affected, at the expression level, by NOR; more specifically, as shown in Figure 3D, expression of *SlNAP2* is significantly reduced in leaves of the *nor* mutant, while it is elevated in the *NOR* overexpressor line *OX-19*, suggesting that NOR acts upstream of *SlNAP2.* On the other hand, we found that expression of *NOR* is enhanced in *SlNAP2-IOE* plants shortly (6 h) after EST treatment (Figure 7B), consistent with a model that places SlNAP2 upstream of *NOR.* Collectively, the available experimental data therefore suggest that NOR and SlNAP2 together form a positively acting regulatory loop whereby the expressional activity of each *NAC* gene is enhanced by the respective other NAC transcription factor. However, we note that unravelling the details of this regulatory interaction require further detailed investigation in the future.

Together, the available data strongly suggest that NAC transcription factors controlling leaf senescence also affect age-dependent senescence (or ripening) of fleshy and non-fleshy fruits, across species. This observation raises a number of interesting questions, including the following: (i) How do NAC TFs exert their specific aging-related functions in photosynthetic leaves compared to those in fruits, i.e., how are the target genes prevalent or specific for leaf senescence selected compared to target genes involved in fruit ripening? (ii) Related to this: do NAC TFs interact with different other transcription factors in leaves *versus* fruits to exert their molecular functions? (iii) To what extent do epigenetic marks affect which genes are primary targets of the senescence-related NACs in leaves *versus* fruits? (iv) In which way has evolution shaped the gene regulatory landscape of age-related NAC TFs in leaves compared to fruits? These questions lead to an even wider perspective which addresses the diversification of NAC functions at the organ, tissue and cellular levels, an aspect not well understood at present. Future research clearly has to address this aspect in more detail.

**Figure 8.**
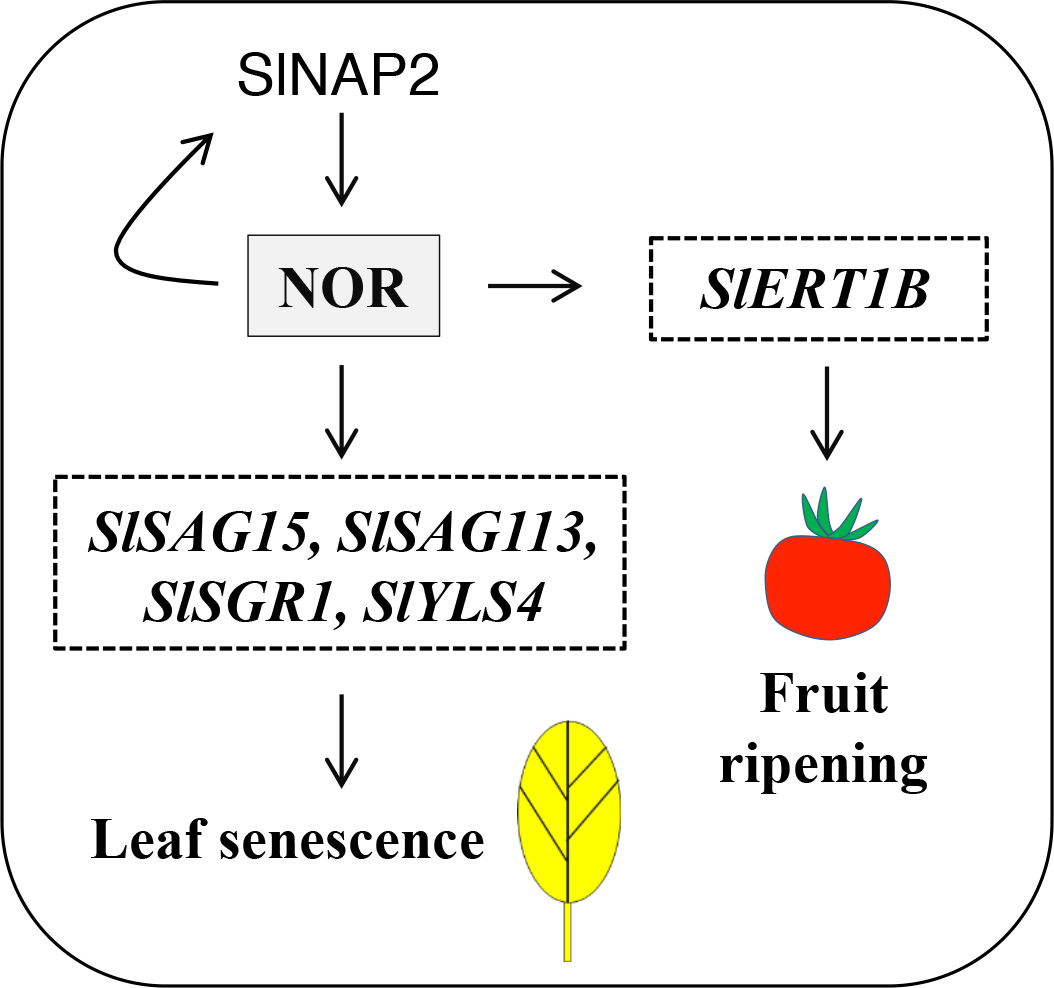
Model for the regulation of leaf senescence by NOR. NOR positively controls leaf senescence in tomato by directly regulating various senescence-associated genes including, besides others, *SlSAG15*, *SlSAG113*, *SlSGR1* and *SlYLS4.* Furthermore, the previously reported NAC transcription factor SlNAP2 (Ma *et al.*, 2018) enhances *NOR* expression by directly binding to its promoter. In addition, NOR enhances *SlNAP2* expression, suggesting a positively acting feed-forward loop involving the two NAC factors. NOR also directly and positively regulates the expression of the fruit ripening-related gene *SlERT1B*, consistent with its well-known role in this process.

## Acknowledgments

We thank Dr. Karin Koehl and her team (Max Planck Institute of Molecular Plant Physiology) for plant care, and Gang-Ping Xue (CSIRO Agriculture and Food, Australia) for performing the binding site selection assays. We thank the University of Potsdam and the Max Planck Institute of Molecular Plant Physiology for supporting our research

## Supplemental Data

Table S1. Oligonucleotide sequences.

Table S2. Data for results shown in heat maps. Table S3. Promoters of NOR target genes.

Figure S1. Selection of *NOR* transgenic lines.

**(A**) *NOR* expression in *NOR* overexpression lines, *OX-L5* and *OX-L19*, compared to WT. Expression was analyzed in 3-week-old seedlings. Data are means ± SD (n = 3). Significant differences from WT are indicated by asterisks (Student’s *t*-test, **: *P* ≤ 0.01). **(B**) Expression of *NOR* in *NOR-IOE* plants. *NOR* expression was analysed by qRT-PCR in *NOR-IOE* plants treated with estradiol (15 μM) for 6 h, or with ethanol (0.15% [v/v]); Mock). Expression of *NOR* after EST treatment is significantly higher than in mock-treated plants. Data are from three biological replicates (Student’s *t*-test; **: *P* ≤ 0.01).

Figure S2. Dark-induced senescence in *ami-NOR* plants.

**(A**) Schematic representation of the *NOR* coding sequence showing the position targeted by the *amiRNA.* Numbers indicate nucleotide positions relative to the *ATG* start codon. A 21-bp sequence (TGTACCATAGTTTGAAGGCTG) was targeted towards the NOR coding sequence. Two transgenic *ami-NOR* lines were selected, namely *ami-L2* and *ami-L35.* Expression was analyzed by end-point PCR amplifying the full-length *NOR* transcript (lower panel). Note the strong downregulation of *NOR* transcript abundance in the *ami-NOR* lines compared to wild type (WT). **(B**) Young leaves detached from 10-week-old WT and *ami-L35* plants after 9 days of dark treatment. Note the less advanced senescence in *ami-L35* compared to WT. **(C**) Chlorophyll content in leaves of WT and *ami-L35* plants before (Control) and after dark incubation for 9 days (Dark), determined using a SPAD meter. Values represent the mean ± SD of three biological replicates each (Student’s *t*-test, *: *P* ≤ 0.05).

Figure S3. Identification of the binding sequences of NOR.

**(A**) Binding activities of NOR toward TaNAC69-selected oligonucleotides. Binding activity is expressed relative to that of S1 (arbitrarily set to 1). The core binding motif is highlighted in red. RBA, relative binding activity. **(B**) Mutational analysis. Mutated S1 motifs (S1m1 ‒ S1m17) and mutated S10 motifs (S10m1 and S10m2) were included in the analysis. For base-substitution analysis, substituted bases in S1 and S16 are shown in lower-case blue letters. Bases inserted are shown in blue and underlined. Values are means of two assays. The data indicate CGTR (5-7N) NACGHMWVH and as high-affinity binding sites of NOR (B = CGT; W = AT; Y = CT; M = AC; H = ACT; V = ACG; N = ACGT). NOR shows more tolerance to mutation of S1 that contains two core motifs, compared to mutations of S10 which has only one core motif.

